# Evidence that many prokaryotic accessory genes are adaptive

**DOI:** 10.64898/2026.03.16.712127

**Authors:** Ying Chen Eyre-Walker, Cara Conradsen, Michiel Vos, Adam Eyre-Walker

## Abstract

Bacterial genomes often contain many genes that are only present in a subset of strains, the so-called accessory genes. Whether these genes are adaptive, neutral or deleterious remains contentious. Here we introduce a simple test to differentiate between these possibilities. If an accessory gene is adaptive then the sequence of the gene should be conserved, and the ratio of non-synonymous to synonymous diversity, *π*_*n*_⁄*π*_*s*_, should be less than one. In contrast, if the gene is neutral or deleterious, selection should not conserve the gene sequence, and *π*_*n*_⁄*π*_*s*_ should equal one. We apply this test to accessory genes in *Escherichia coli* and *Staphylococcus aureus*, two highly divergent bacterial species with a large and a small pangenome respectively. We find *π*_*n*_⁄*π*_*s*_<1 for genes at all frequencies in both species demonstrating that many are adaptive. We estimate that at least 75% of all the accessory genes are maintained by selection in the two species that we have analysed, equating to thousands of adaptive accessory genes in both species, a substantial increase on previous estimates.

**Significance statement:** Many bacterial genes are found in only a subset of the strains of a bacterial species. These are referred to as accessory genes. Whether, accessory genes are adaptive, allowing certain strains to inhabit particular niches, remains a highly contentious question. In this paper we introduce a simple test of whether accessory genes are adaptive and show that >75% of them are, in two highly divergent bacterial species. This radically alters our view of this type of genetic variation and bacterial evolution.

## Introduction

One of the most astonishing discoveries from the genomic era has been the observation that bacterial genomes of the same species often have very different gene repertoires; for example, two *Escherichia coli* genomes typically only share 70% of their genes (1, 2). Genes that are present in some, but not all strains are referred to as accessory genes (AGs), whilst those that are present in all strains are known as core genes; together the accessory and core genes form the pangenome.

Why do AGs exist? There are two main hypotheses. First, it has been suggested that AGs are neutral or deleterious genetic variation caused by the introduction of new genes by selfish genetic elements or by the cell itself through the process of natural transformation (3). They may also represent cases in which a previously core gene has been lost by deletion in a subset of strains, the deletion being either neutral or deleterious (3). Second, it has been suggested that AGs may represent genetic variation that is beneficial to the bacterium in some of the niches that it inhabits (4).

Which of these processes dominates the evolution of AGs remains debated but unresolved (3-6). Early studies sought to differentiate between the explanations by fitting evolutionary models to the distribution of genes across strains (7-10), the so-called gene frequency distribution (GFD) or spectrum (GFS). The GFS is typically U-shaped, with most genes occurring in either very few strains (known as the “cloud”), or in all or nearly all strains (“core” or “soft-core”), while only a small minority are found at intermediate frequencies (“shell”) (9). These early studies found that the GFS was consistent with a model in which AGs were neutral, so long as there were a few categories of genes with different rates of gain and loss (7-10), which seems realistic (11). Whether models of selection can also explain the GFS within a species has not been investigated.

More recent analyses have taken a functional approach to testing for adaptation. Goyal (12) used metabolic models to predict whether AGs produced additional metabolites that might be beneficial across 96 species; this was indeed the case for 80% of the species they considered. Furthermore, randomising AGs across strains produced significantly fewer metabolites, even when the randomisation kept operons intact. Consistent with these co-dependencies, Beavan et al. (13) found that the presence of some genes was predictable based on other genes present in the genome. Taking a different approach, Douglas and Shapiro (14) found that for many categories of AGs the likelihood of being functional, as opposed to being a pseudogene, was increased by the absence of a functionally similar gene in the genome, consistent with some AGs being adaptive. They estimated that ∼20% of AGs in their sample were maintained by selection.

Note there have also been studies that have tested for adaptation in the evolution of gene content between species (15, 16). These analyses often suggest a role for adaptation; however, it is possible for most of the variation within a species to be neutral or deleterious, and all differences between species to be a consequence of adaptive evolution. Therefore, these studies do not resolve whether AGs segregating within a species are adaptive, neutral or deleterious.

Whilst functional analyses have produced limited evidence of adaptation, the methods are complex and hence potentially subject to bias. Here, we introduce a new and simple method to test whether AGs are adaptive. If AGs are adaptive, then there should be selection to conserve the sequence; hence the ratio of non-synonymous to synonymous nucleotide diversity (*π*_*n*_⁄*π*_*s*_)should be less than one, because many non-synonymous mutations will be deleterious and hence removed from the population by natural selection. In other words, we infer that a gene is advantageous from the evidence of negative selection against changes to the sequence of the gene. In contrast, if an AG is neutral or deleterious there will be little or no selection to maintain its function and hence the sequence, and *π*_*n*_⁄*π*_*s*_ should be equal to one. Here we apply this test to the AGs of two bacterial species, *Escherichia coli* and *Staphylococcus aureus*.

## Results

We applied our new test of whether AGs are adaptive to the two highly divergent and well sampled bacterial species *Escherichia coli* and *Staphylococcus aureus*. For each species we obtained a pangenome constructed from 500 strains from the PanX database (17). Although both show the classic U-shape gene frequency distribution (Figure S1), *E. coli* has a relatively open pangenome with 28,297 genes, whilst *S. aureus* has a relatively closed pangenome of only 6,139 genes. Of these, 97% and 88% are accessory (i.e. they are not found in every strain) respectively. Because our test is based on estimating AG nucleotide diversity, it can only be applied directly to AGs present in two or more strains (although see section below) and 66% and 83% of the AGs satisfy this criterium. Hence, we can apply our test to a substantial fraction of the AGs present in the samples from the two species.

In both species average *π*_*n*_⁄*π*_*s*_ is substantially and significantly less than one in AGs, consistent with many AGs being adaptive and maintained by selection (Figure 1A). This pattern is consistent across AGs in the lowest frequency class (2 strains), at low (3-50 strains), intermediate (51 to 450 strains) and high (451 to 499) frequencies (P<0.001 in all cases) (Figure 1A, Table S1). In fact, *π*_*n*_⁄*π*_*s*_ is significantly less than 1 (at P<0.05) for many individual gene frequency categories – in all 95 gene frequency categories that have ten or more genes in *E. coli* and in 37 out 39 gene frequency categories in *S. aureus*.

**Figure 1.**
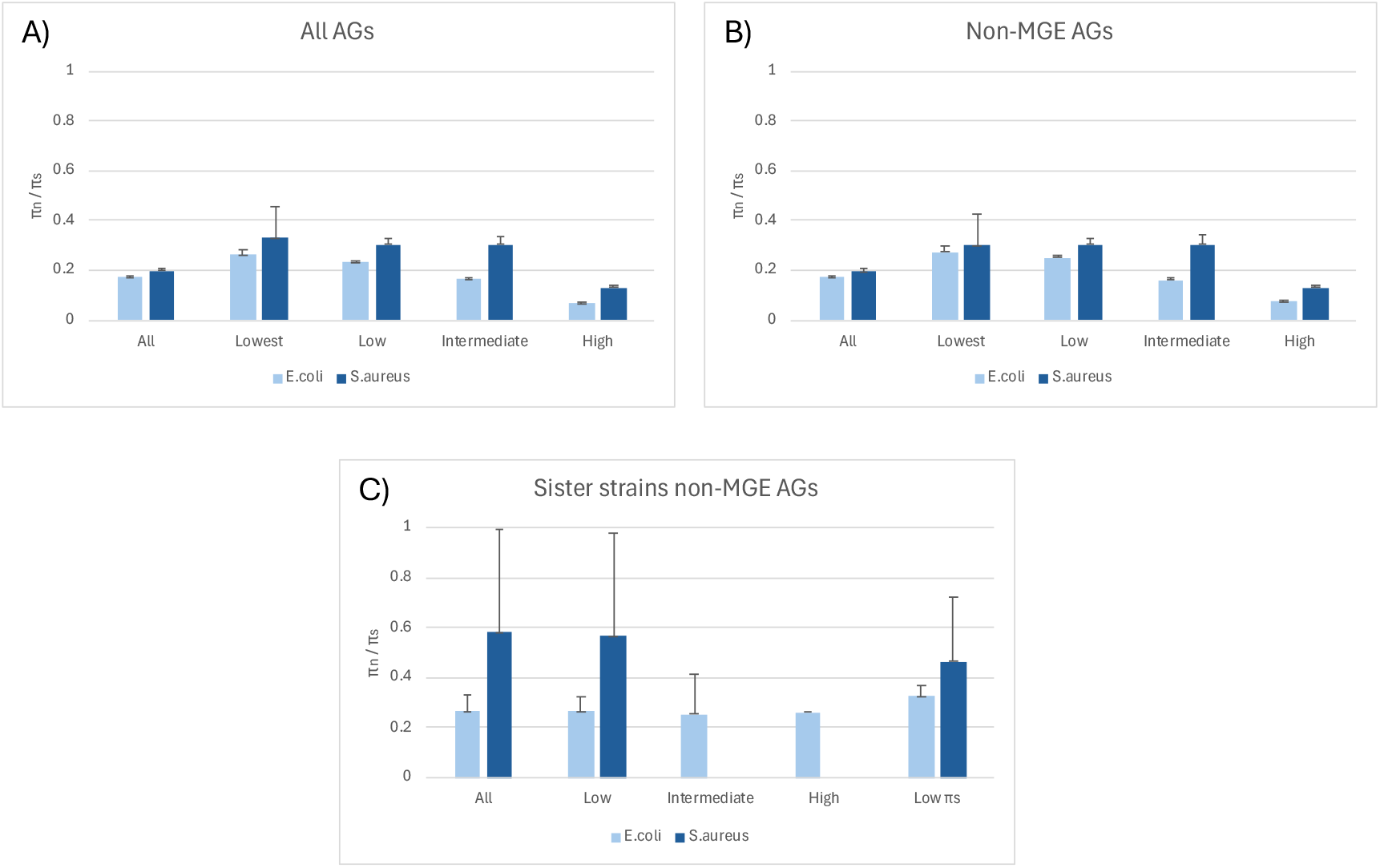
The average *π*_*n*_⁄*π*_*s*_ for different categories of AGs (A) all genes, (B) all non-MGE related genes, and C) the divergence between sister strains for all non-MGE related genes. The average *π*_*n*_⁄*π*_*s*_ is shown for all genes, those at the lowest (2 strains), low (2-50 strains), intermediate (51-450 strains) and high (451-499 strains) frequencies, except, for the sister strain non-MGE plot, where we combine the lowest and low frequency categories due to insufficient data in *S. aureus*; *π*_*n*_⁄*π*_*s*_ is undefined for the intermediate and high frequency classes in *S. aureus* because *π*_*s*_= 0 for the point estimate or some of the bootstrap replicates. The upper 95% confidence intervals are shown, as determined from 10,000 bootstrap replicates.

### Mobile genetic elements

The fact that the average *π*_*n*_⁄*π*_*s*_ is less than one is consistent with many AGs being adaptive. However, we need to be cautious. A substantial number of AGs are derived from mobile genetic elements (MGEs)(4, 18-20) and the sequence of an MGE might be conserved because only MGEs with intact genes can mobilise; such AGs might not be beneficial and may even be detrimental to the host. To investigate this, we used a variety of bioinformatic tools and MGE databases to annotate each genome (see Methods). We find that 14% of AGs are related to MGE maintenance and propagation in both species; we refer to these as MGE genes. As expected, *π*_*n*_⁄*π*_*s*_<1 in MGE genes (*E. coliπ*_*n*_⁄*π*_*s*_=0.176 (0.005); *S. aureusπ*_*n*_⁄*π*_*s*_ = 0.248 (0.019)). However, if we remove MGE genes from the analysis we find that *π*_*n*_⁄*π*_*s*_is still significantly less than one overall, and for each frequency group, consistent with many AGs being adaptive (Figure 1B).

### Multiple introductions

We also need to be cautious because some AGs may have been introduced to a pangenome multiple times. There is a possibility that the observed sequence conservation could be due to selection in the donor species where the AG could be core, rather than in the recipient where it is accessory. To investigate whether multiple introductions affect our results, we restricted our analysis to the divergence between pairs of sister taxa on a phylogenetic tree constructed from the core genome. For example, a gene might be present in strains A, B, C and D, where strains A and B, and C and D are sister strains; we only consider the divergence in the gene between A and B, and between C and D, but not between A and C etc. Genes shared between sister taxa are likely to be the result of vertical inheritance after a single introduction to the ancestor of the strain pair (even if the gene has been introduced multiple times across the phylogeny). We find that *π*_*n*_⁄*π*_*s*_ is significantly less than one for all such AGs no matter their frequency in *E. coli*; in *S. aureusπ*_*n*_⁄*π*_*s*_ is significantly less than one for all genes and for rare AGs; however, we have too little data to estimate *π*_*n*_⁄*π*_*s*_ in genes at intermediate and high frequency.

As an additional check, we restricted our analysis to genes with low synonymous diversity. These are likely to be the product of a single introduction for two reasons. First, because they have low *π*_*s*_ they are more likely to be recently, and therefore singly, introduced. Second, genes that have been introduced multiple times will tend to have high diversity because they will be sampled from the diversity of the donor. Since selecting genes by their *π*_*s*_ value will increase the estimate of *π*_*n*_⁄*π*_*s*_ by sampling error, we divided the gene into odd and even codons using *π*_*s*_(odd) to select the genes and *π*_*s*_(even) to estimate *π*_*n*_⁄*π*_*s*_ (21). We restrict our analysis to non-MGE related genes >200bp in length to increase the power of the analysis and to those with *π*_*s*_(odd) = 0, the lowest threshold we can select. We find that mean *π*_*s*_(even) = 0.002 and 0.0007 in *E. coli* and *S. aureus* respectively; these are more than an order of magnitude lower than the average *π*_*s*_ for core genes - 0.040 and 0.022 in *E. coli* and *S. aureus* respectively – and lower than *π*_*s*_ found in most bacterial species (22), suggesting that the genes are unlikely the consequence of multiple introductions. We find that *π*_*n*_⁄*π*_*s*_ is significantly less than one in both species even in these genes with very low *π*_*s*_ values (Figure 1C; Table S1).

### Proportion of adaptive AGs

It is possible to estimate the proportion of AGs that are adaptive using a simple model. Let us assume that a proportion α of AGs are MGE-related and that a proportion β of the non-MGE related genes are adaptive, with the other non-MGE related genes being either neutral or deleterious. Under this model average *π*_*n*_⁄*π*_*s*_ is

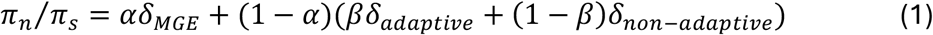

where *δ*_*MGE*_, δ_*adaptive*,_ and *δ*_*non*-*adaptive*,_are the expected values of *π*_*n*_⁄*π*_*s*_ for MGEs, adaptive non-MGEs and non-adaptive non-MGEs respectively. Equation 1 can be rearranged to yield an estimate of β, the proportion of non-MGE genes that are adaptive. If we assume that δ_*MGE*_ is as we observe in our data, that δ_*adaptive*,_ is the value of *π*_*n*_⁄*π*_*s*_ found in highest frequency AGs; note that *π*_*n*_⁄*π*_*s*_ is negatively correlated to gene frequency (see below) and hence this a conservative lower estimate for δ_*adaptive*_. If we further assume that δ_*non*-*adaptive*,_ = 1, as we expect under our model, then we estimate that 76% and 78% of all the AGs that we have analysed are adaptive in *E. coli* and *S*. *aureus*, respectively, given that 14% of AGs are classified as MGE genes. This equates to 13,800 and 3,500 genes in the two species. If we assume that AGs are less conserved than core genes, then our estimates will increase and could potentially be 86% - i.e. all non-MGE genes.

### Singleton AGs

We estimate that more than 75% of all AGs present in 2 strains or more are adaptive. However, a large proportion of AGs are found in just one strain in each species (34% in *E. coli* and 17% in *S. aureus*). To investigate whether these might also be selectively constrained and hence adaptive we considered the relationship the *π*_*n*_⁄*π*_*s*_ and gene frequency. We find that *π*_*n*_⁄*π*_*s*_ is negatively correlated to gene frequency in both species (All genes: *E. coli* r=-0.15, p=0.002; *S. aureus* r=0.18, p=0.003, non-MGE related genes: *E. coli* r=-0.16, p=0.001; *S. aureus* r=-0.17, p=0.006)(Figure 2), however the slope is shallow and there is no evidence that the relationship is non-linear even if we consider the low frequency categories (Figure 2B and 2C). From the linear regression we predict *π*_*n*_⁄*π*_*s*_ to be 0.24 and 0.35 for non-MGE genes present in a single strain for the *E. coli* and *S. aureus* respectively, essentially identical to the values for genes present in 2 strains. Hence, we infer that most genes present in a single strain are also adaptive.

**Figure 2.**
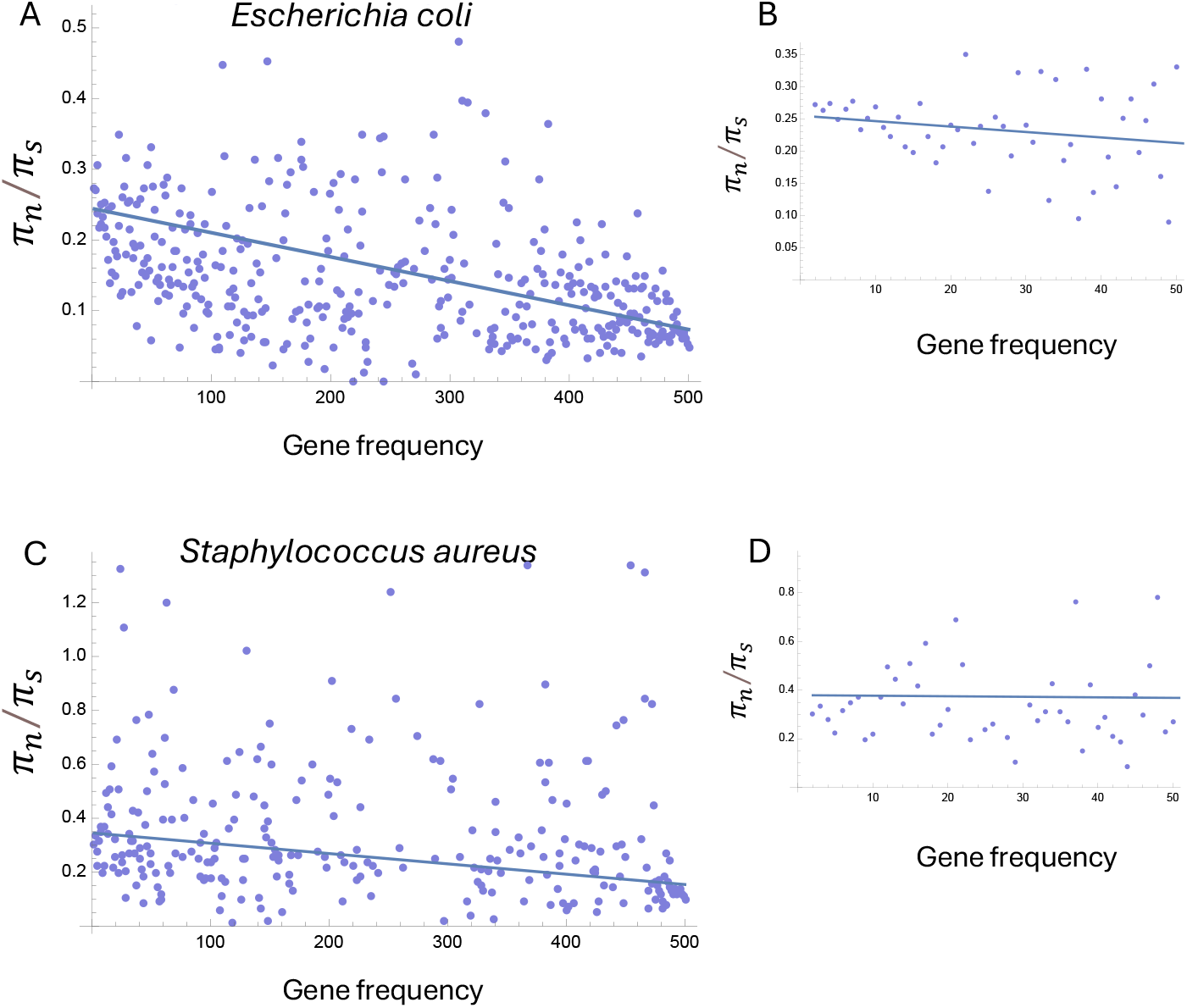
The relationship between *π*_*n*_⁄*π*_*s*_ and gene frequency for non-MGE related genes for *E. coli* (A and B) and *S. aureus* (C and D). The line of best fit is shown; in each case this has a slope which is significantly less than zero (p<0.01). B and D show the relationship for the lowest frequency genes.

## Discussion

We have demonstrated that many AGs are subject to substantial selective constraint in their sequence. This is consistent with many AGs being adaptive – natural selection has maintained their sequence because they are advantageous to their host. On this basis we estimate that at least 75% of the AGs that we have analysed in both species are adaptive. However, there are several caveats.

First, it is essential that we only consider genes that are not part of an MGE, since the sequence of an MGE might be conserved for reasons other than its effects on host fitness. Although, we have filtered our data against a broad range of MGE databases, this filtering is inherently incomplete since new MGEs are being continuously discovered, and no bioinformatic tool captures all elements with perfect sensitivity (23). Using equation 1 we can investigate the consequence of incomplete MGE annotation. If we assume that *δ*_*MGE*_ is the same for identified and non-identified MGE related genes, and that *δ*_*adaptive*,_ is equal to *π*_*n*_⁄*π*_*s*_ in the highest frequency AGs, as above, then we find that the proportion of AGs inferred to be adaptive decreases if we increase the proportion of genes that are MGE; although note that the proportion of genes inferred to be adaptive amongst the non-MGE genes is hardly affected (Table 1). However, we would still infer that the majority of AGs are adaptive, i.e. beneficial to the host, even if there are three times as many AGs as we have detected (Table 1).

**Table 1.**
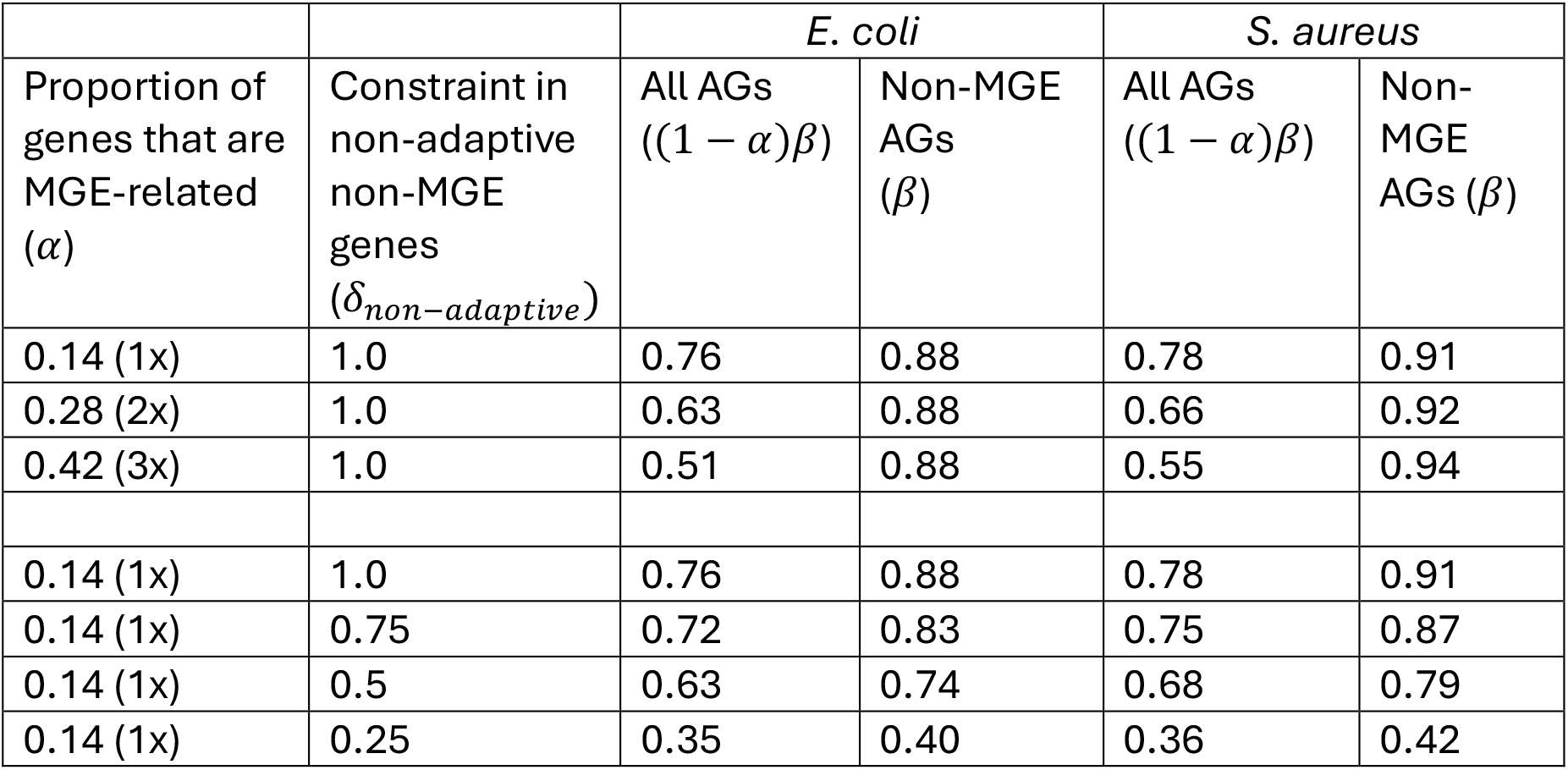
The proportion of all accessory genes, and non-MGE related accessory genes that are estimated to be adaptive using equation 1 as a function of the proportion of genes that are MGE-related and the constraint upon non-adaptive non-MGE genes.

| Proportion of genes that are MGE-related ($\alpha$) | Constraint in non-adaptive non-MGE genes ($\delta_{non-adaptive}$) | *E. coli* | | *S. aureus* | |
| --- | --- | --- | --- | --- | --- |
| | | All AGs $((1 - \alpha)\beta)$ | Non-MGE AGs ($\beta$) | All AGs $((1 - \alpha)\beta)$ | Non-MGE AGs ($\beta$) |
| 0.14 (1x) | 1.0 | 0.76 | 0.88 | 0.78 | 0.91 |
| 0.28 (2x) | 1.0 | 0.63 | 0.88 | 0.66 | 0.92 |
| 0.42 (3x) | 1.0 | 0.51 | 0.88 | 0.55 | 0.94 |
| 0.14 (1x) | 1.0 | 0.76 | 0.88 | 0.78 | 0.91 |
| 0.14 (1x) | 0.75 | 0.72 | 0.83 | 0.75 | 0.87 |
| 0.14 (1x) | 0.5 | 0.63 | 0.74 | 0.68 | 0.79 |
| 0.14 (1x) | 0.25 | 0.35 | 0.40 | 0.36 | 0.42 |

Second, we need to restrict our analysis to genes that have been introduced a single time into our focal species, or if the gene has been introduced multiple times, we need to consider the evolution amongst the descendants of each introduction. To investigate this, we performed two analyses. In the first we restricted the analysis to the evolution between sister strains, which are likely the consequence of a single introduction. In the second analysis we considered AGs with very low synonymous diversity, a proxy for genes that had been recently introduced and hence likely to be the product of a single introduction. In both tests in both species, we find that *π*_*n*_⁄*π*_*s*_ is significantly less than one, consistent with the non-MGE AGs being adaptive in the species we are considering.

Third, there is also a potential neutral explanation for our results. It is possible that AGs are neutral or deleterious, but their sequence is constrained by selection to avoid becoming toxic, through protein misfolding for example (24). Recent experimental work in *E. coli* and *S. cerevisiae* suggests that the toxic effects of protein misfolding exert little selective pressure on genes beyond reducing the concentration of the protein (25, 26). However, to investigate whether this type of selection is likely to affect our results we reduced the value of *δ*_*non-adaptive*,_ in equation 1 from its assumed value of 1.0, to 0.75, 0.5 and 0.25; these values would imply that 25%, 50% and 75% of non-synonymous in non-adaptive non-MGE genes generate toxic protein products and are selected against. We find that our estimates of the proportion of genes that are adaptive is affected by assuming that a proportion of mutations are selected against; however, we estimate that the majority of AGs are adaptive even if 50% of amino acid mutations had these effects (Table 1).

It may seem surprising that so many AGs in our sample are adaptive given that we might expect the introduction of a new gene to be deleterious because of the energetic costs in terms of DNA replication, and if the gene is expressed, the costs of transcription, translation and epistatic fitness interactions (27). There are two potential resolutions of this conundrum. First, most new genes introduced into a bacterium may be deleterious and removed by natural selection, with only those that are strongly beneficial overcoming these costs and spreading to a frequency that they are observed in a modest sample of strains, of the sort that we have analysed. Second, it is possible that many of the genes that are acquired by HGT are advantageous. The probability that a gene is transferred to a new host is likely to depend upon its frequency in the various potential donor species; hence genes that are beneficial to many potential hosts might be more likely to be those that are transferred frequently.

We have estimated that >75% of AGs in our sample are adaptive; this is a substantial increase on the previous estimate of 20% derived by a different method (14). However, our estimate is not directly comparable to that of Douglas and Shapiro since they included genes present in a single strain in their analysis whereas we have not, and these comprise a substantial fraction of AGs. Nevertheless, as we argue above the relationship between *π*_*n*_⁄*π*_*s*_ and AG frequency suggests that genes present in a single strain are likely under constraint and hence adaptive.

Our estimate only pertains to the genes at a sufficient frequency to be sampled; approximately those with a frequency of 1/500 or above. If we sequence more strains, then we will discover more genes at lower frequencies, and it is likely that an increasing proportion of these will be neutral or deleterious. To estimate the proportion of all genes in the pangenome of a species we need to know the underlying gene frequency distribution (not the frequency in the sample, but in the whole species) and the relationship between *π*_*n*_⁄*π*_*s*_ and AG frequency. These are likely to be difficult to estimate.

The nature of selection acting upon AGs remains far from clear. There are several potential non-exclusive models. First, AGs might be retained through some form of balancing selection. This might take the form of many spatially separated niches, each of which requires a different set of genes (4). It might also involve temporally changing environments. There is a growing appreciation that selection fluctuates through time for many organisms (28); in some circumstances these fluctuations are predictable - for example the changing of the seasons (28, 29). But in other situations, the environment might change erratically (30-32). It is also possible that there is frequency dependent selection (33) or even heterozygous advantage where the chromosome and a plasmid share a locus (25). Finally, it might be that many AGs are weakly beneficial over many environments such that they are subject to directional selection (34). We currently do not have sufficient data to differentiate between these models, and all may affect AGs.

Genes associated with MGEs also show significant constraint on their sequence (*E. coliπ*_*n*_⁄*π*_*s*_ = 0.176; *S. aureusπ*_*n*_⁄*π*_*s*_ = 0.248), similar to that on non-MGE AGs (*E. coliπ*_*n*_⁄*π*_*s*_ = 0.171; *S. aureusπ*_*n*_⁄*π*_*s*_ = 0.193). Selection for MGE sequence conservation may come from three non-mutually exclusive sources: selection to maintain the ability to mobilise the element, selection favouring MGE genes that are beneficial to the host, and selection against toxic non-synonymous mutations.

We find that *π*_*n*_⁄*π*_*s*_ is negatively correlated to gene frequency in both species. This is consistent with several different models. It may reflect the fact that some of the genes at low frequency are neutral or deleterious. But it may also be due to consistently lower levels of selective constraint on low frequency genes. Finally, it may be due to differences in the effective population size: low frequency genes are present in fewer copies in the population and hence have a lower effective population size, with less effective selection against deleterious genetic variation within the sequence. It is currently difficult to differentiate between these possibilities.

In summary, we have investigated whether accessory genes are adaptive in two divergent bacterial species. We find strong evidence that more than 75% of the non-MGE related genes are conserved in their protein coding sequence and hence are adaptive.

## Materials and methods

We downloaded the gene alignments for the pangenomes of *Escherichia coli* and *Staphylococcus aureus* from the PanX database (pangenome.org) (17). These were constructed from 500 genome sequences each. The bacteria were selected because they are highly divergent from each other, one being gram positive and the other gram negative; they also have different pangenomes sizes with the pangenome of *E. coli* containing almost 4x as many genes as *S. aureus*. PanX provides alignments for all genes in the pangenome.

The data were filtered to remove a few genes that contained duplicate genes from the same strain. For genes present in 2 or more strains we estimated the number of synonymous and non-synonymous differences per site, *d*_*s*_ and *d*_*n*_ respectively, using the method of Li et al. (35) implemented with the *kaks* function from the R package Seqinr (36) between all pairs of strains for which the gene was present. The non-synonymous (*π*_*n*_) and synonymous (*π*_*s*_) nucleotide diversities were then estimated as the average of *d*_*n*_ and *d*_*s*_ across all pairs of strains with the gene. We found that some genes showed very high diversity and visual inspection suggested that not all sequences in the alignment were true orthologs. We therefore removed the 10% of genes with the highest *π*_*s*_ values. We estimated the average value of *π*_*n*_⁄*π*_*s*_ for a group of genes by summing the values of *π*_*n*_ and *π*_*s*_ across genes; we did this for two reasons. First, for many genes *π*_*s*_ = 0 and hence *π*_*n*_⁄*π*_*s*_ is undefined, and second by aggregating data in this way we minimise the bias inherent in estimating ratios.

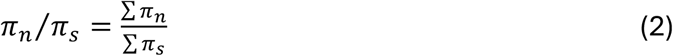

We derived the confidence intervals on this estimate by bootstrapping the data by gene 10,000 times.

We restricted some of our analysis to genes present in sister strains. Sister strains were inferred from a phylogenetic tree constructed from the core genes; this tree was downloaded from the PanX website (pangenome.org)(17).

For some analyses we removed genes associated with Mobile Genetic Elements (MGEs). These were identified in the following manner. We downloaded the accession names for the *E. coli* and *S. aureus* PanX pangenomes to map loci to plasmid or chromosomal scaffolds using NCBI genome annotations. From each gene alignment, the longest and most frequent sequence was used as the representative sequence for annotation. Sequences were bulk annotated using Bakta v1.8.1, via the bakta_proteins function, which includes assignment of COG categories and IS elements via ISFinder (37).

MGEs genes were distinguished from bacterial housekeeping and other accessory genes using MGEfams [v0.5] (38) and mobilOG-db [Beatrix 1.6 v1](23), manually curated databases that exclude accessory genes such as antimicrobial resistance, metal resistance, and virulence factors. To complement these databases, further functional annotation was carried out by performing BLASTp on the loci against MGE-related databases: TnCentral [v. 2](39), PHASTEST [Prophage database; Dec 22, 2020](40), and PLSDB [v. 2024_05_31_v2](41). Candidate hits were first filtered by selecting, for each query, the entry with the maximum bit score, followed by the lowest E-value, highest percent identity, longest alignment length, and then longest query length. Finally candidate hits were retained if they met the following thresholds: bitscore ≥ 60, sequence identity ≥80%, E-value ≤1e-5, and query alignment length ≥70 amino acids.

Annotated loci were then classified using a stepwise pipeline. Insertion sequences were identified by regular expression matching against ISFinder-associated keywords, while hypothetical or uncharacterized proteins were classified based on keywords (e.g., ‘hypothetical,’ ‘DUF,’ ‘ORF’) or assignment to COG category S. MGEs were defined by hits to mobOG, MGEfams, TnCentral or COG category X. Phage and plasmid associated loci were classified using PHASTER and PLSDB hits, respectively, while excluding defence and cargo genes that were identified using CARD, VFDB, BacMet, mobOG, or COG categories V and P. Remaining loci lacking database support were assigned based on residual COG assignments and contextual keywords. Based on these analyses we divided our genes into MGE-related and non-MGE related genes.

All code will be made available at Zenodo.

## Acknowledgements

We have grateful to the Leverhulme Trust for funding (grant number RPG-2024-212) and to James McInerney and Craig Maclean for helpful discussion and comments on the manuscript.

## Supplementary figures

**Figure S1.**
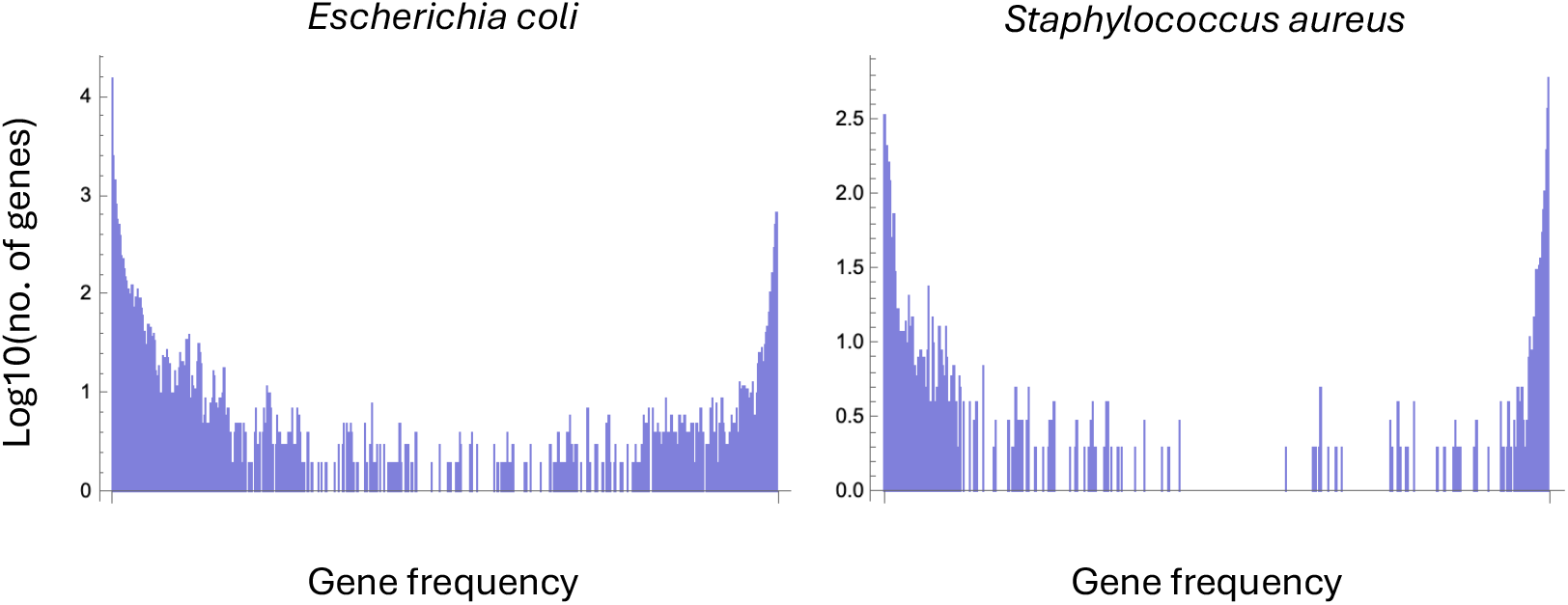
The gene frequency spectra for the pangenomes of *Escherichia coli* and *Staphylococcus aureus*.

**Table S1.**
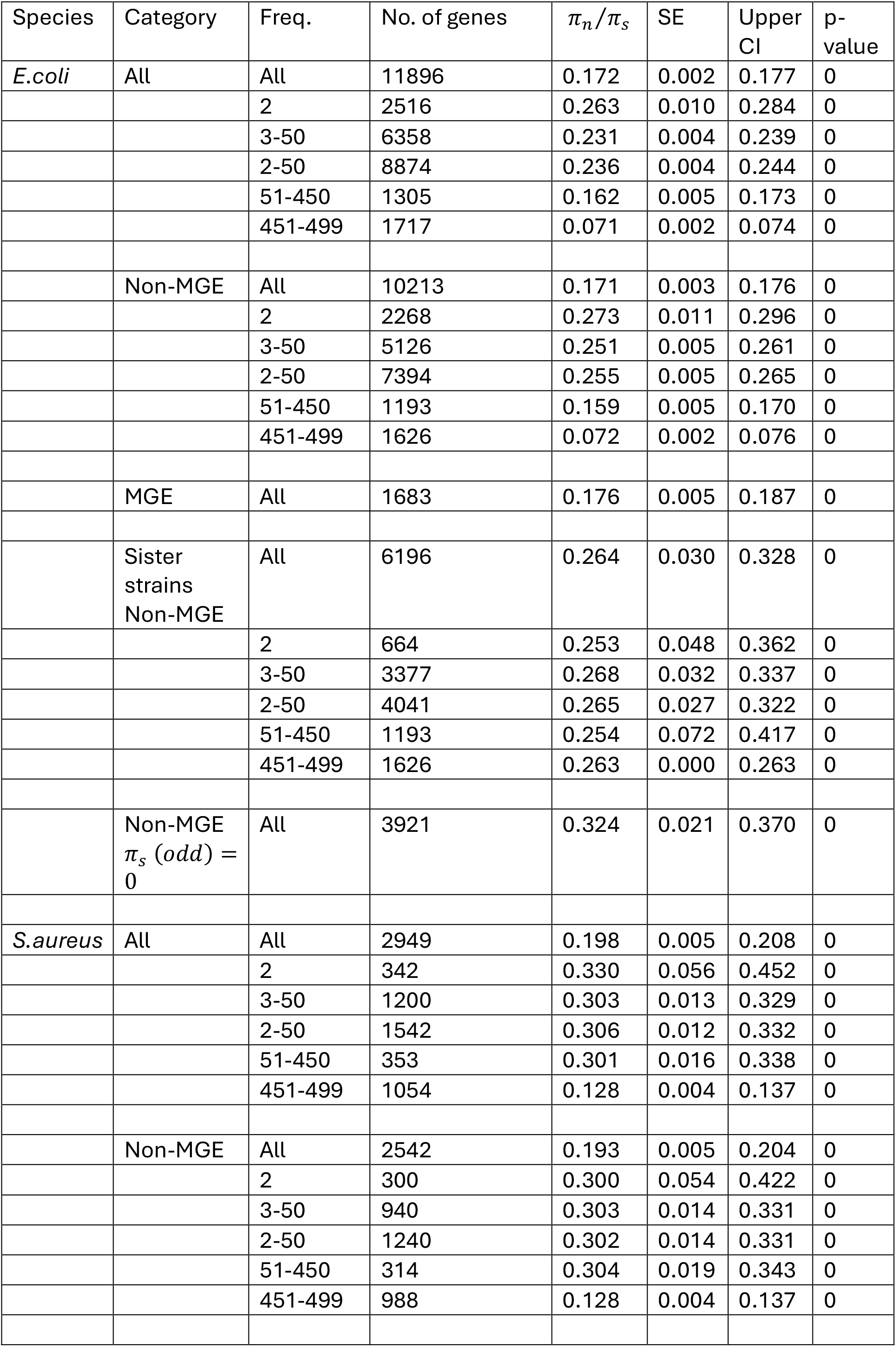
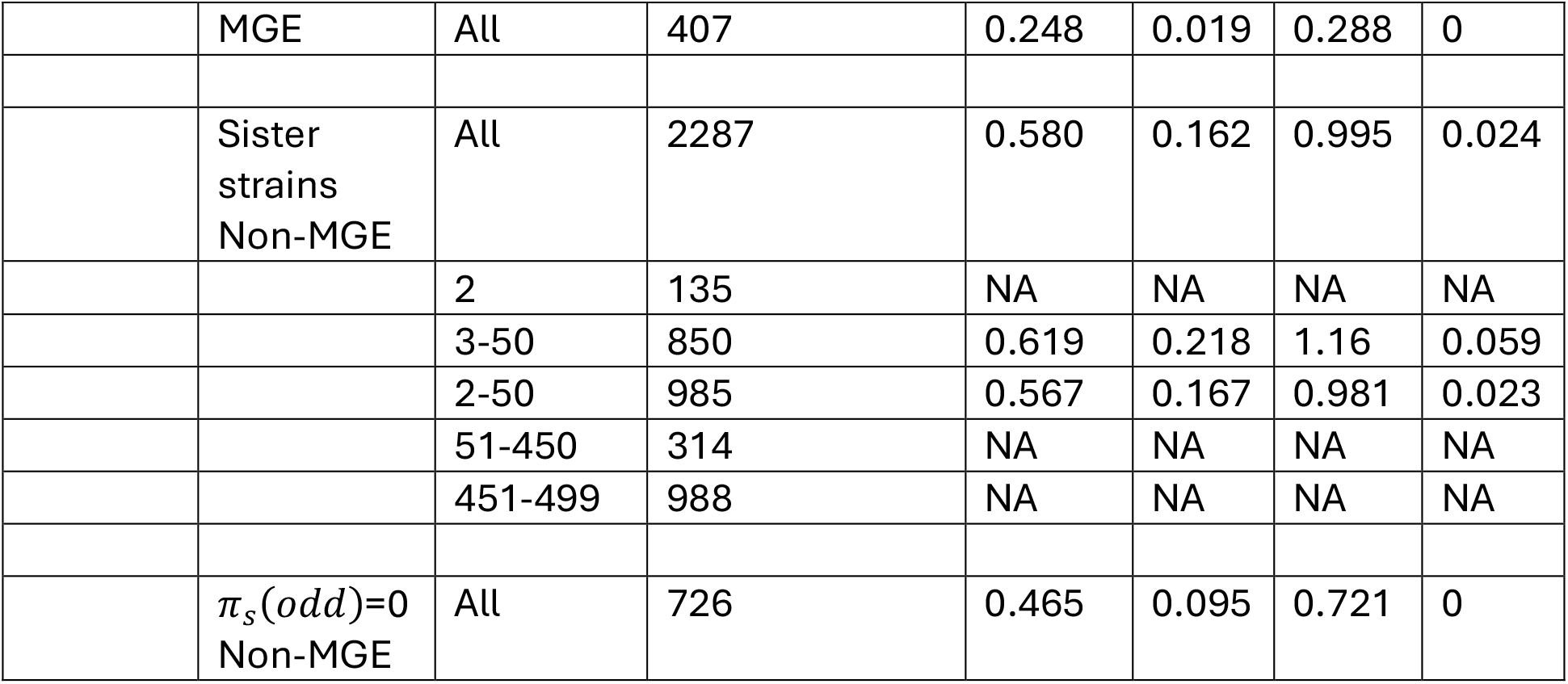
Patterns of selective constraint for different categories of accessory genes. The table reports *π*_*n*_⁄*π*_*s*_, its standard error derived by bootstrapping the data 1000 times, the upper 95% confidence interval and the p-value, the proportion of bootstrap replicates for which *π*_*n*_⁄*π*_*s*_ ≥ 1. Values are given for partitions of the data based on by MGE status and lineage association, and (ii) pangenome frequency (e.g. 2-50 are the results for genes with frequencies of 2 out of 500 to 50 out of 500 strains). *π*_*n*_⁄*π*_*s*_ is undefined or the CI cannot be calculated for some frequency categories in the non-MGE related genes in sister strains analysis in *S*.*aureus* because *π*_*s*_=0 in the point estimate or some of the bootstrap replicates; these are marked as NA.

## Notes

### Competing Interest Statement

The authors have declared no competing interest.

### Summary of Updates

The revision contains several advances. We now consider genes present in a single strain, inferring that most of these are also likely adaptive. We also consider quantitatively whether selection against toxic folding errors will affect our results - we find that I will not unless these effects are very common.

